# A device for exploring the full angular excitation space - Can more angular projections improve determination of a molecules 3D-orientation in the presence of noise?

**DOI:** 10.1101/2020.03.11.986067

**Authors:** Dominik Pfennig, Andreas Albrecht, Julia Nowak, Peter Jomo Walla

**Affiliations:** Institute of Physical and Theoretical Chemistry, Department of Biophysical Chemistry, Technische Universität Braunschweig, Gaußstraße 17, 38106 Braunschweig, Germany

## Abstract

In the past, different methods have been presented to determine the 3D orientation of single molecules in a microscopic set-up by excitation polarization modulation. Using linearly polarized illumination from different directions and thereby measuring different 2D projections enables reconstructing the full 3D orientation. Theoretically, two projections suffice for a full 3D orientation determination if the intensities are properly calibrated. If they are not, a third projection will enable unambiguous orientation measurements. The question arises if three projections already contain the maximum information on the 3D orientation when also considering the limited number of available photons and shot noise in an experiment, or if detecting more projections or even continuously changing the projection direction during a measurement provides more information with an identical number of available photons. To answer this principle question, we constructed a simple device allowing for exploring any projection direction available with a particular microscope objective and tested several different excitation modulation schemes using simulated as well as experimental single molecule data. We found that three different projections in fact already do provide the maximum information also for noisy data. Our results do not indicate a significant improvement in angular precision in comparison to three projections, both when increasing the number of projections and when modulating the projection direction and polarization simultaneously during the measurement.

In fluorescence microscopy polarized illumination from different directions enables the determination of the 3D orientation of single molecules by combining the 2D information of different projection directions. Ambiguities that emerge when using only two projections can be eliminated using a third projection. In a systematic study we show that – also considering the limited number of available photons and shot noise in an experiment – three projection directions already contain the maximum information on the 3D orientation. Our results do not indicate a significant improvement in angular precision in comparison to three projections, both when increasing the number of projections and when modulating the projection direction and polarization simultaneously during the measurement.

## 1. Introduction

In fluorescence microscopy several techniques have been developed by which it is possible to reconstruct the spatial orientation of transition dipole moments of fluorescence dyes. Since both the excitation probability and the fluorescence intensity of fluorescence dye molecules are intrinsically anisotropic, orientation dependent fluorescence signals can be obtained by both modifying the excitation or the detection light path. Annular, [1] radially polarized, [2] defocused [3] or multidirectional linearly-polarized [4] illumination have all been utilized to determine molecular orientation, while polarizers, beam splitters and phase masks in the detection path have also been used to the same end. [5–10] In addition to pure orientation information, several groups have recently started to explore the use of polarization to separate structures at subdiffraction distances. [11–16]

In a recent publication [17] we used a method comparable to the one presented by Prummer et al., [4] using linearly polarized light from two different illumination directions (2dir for short, see Fig. 1a) to determine the three-dimensional orientation. To achieve different illumination directions we laterally shifted the excitation light beam before it entered the microscope’s objective using two antiparallel wedge prisms. The objective lens consequently refracts the light away from the optical axis resulting in a tilted illumination light beam (tilt angle *β*). To realize more than two illumination directions, we now extended our previous approach by constructing a rotating holder for the wedge prisms. Changing the position at which the light enters the objective by rotating this holder (rotation angle *ψ*) changes the tilt direction accordingly. For each illumination direction the projection of the orientation of an emitter’s transition dipole moment onto a plane perpendicular to said direction can be determined using polarization microscopy (compare Ref. [18–20]). When more than one direction is used the full three-dimensional orientation in the sample can be reconstructed from the obtained projections.

**Fig. 1.**
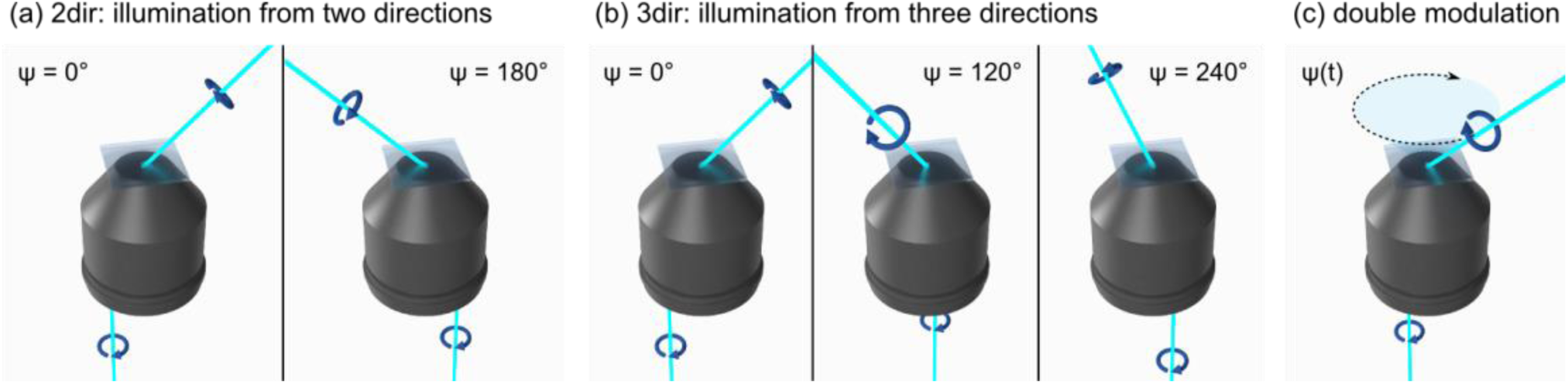
Illumination directions used for 2dir (a) and 3dir (b) simulations and experiments as well as an illustration of the beam’s rotation with the double modulation method (c). The rotation of the illumination beam around the optical axis which directly corresponds to the rotation of the wedge prisms is described by the angle Ψ. The lateral displacement of the beam before entering the objective is exaggerated.

In theory, two projections (Fig. 1a) provide sufficient information about the orientation. Fig. 2 is illustrating this with special easily understood examples: Even if the amplitudes are equal for two molecules, the phases are different (i). If, on the other hand, the phases are equal, the amplitudes will be different (ii). There is no case for which the amplitude and phase is identical for two molecules for both directions. However, for molecules oriented in the plane spanned by the two directions of projection precise knowledge about the absolute brightness in both projections is necessary for an exact and unambiguous orientation reconstruction. Pure phase information does not suffice, as all molecules in this plane have the same pair of phases, and even a relative comparison of the amplitudes is ambiguous (iii). Therefore, in experiments a brightness calibration is vital for unambiguous orientation reconstruction if only two projections are used. Additionally, in experiments data is distorted by background and shot noise as well as blinking of the dyes, increasing the probability of a false orientation reconstruction for molecules oriented close to the plane spanned by the projection directions.

**Fig. 2.**
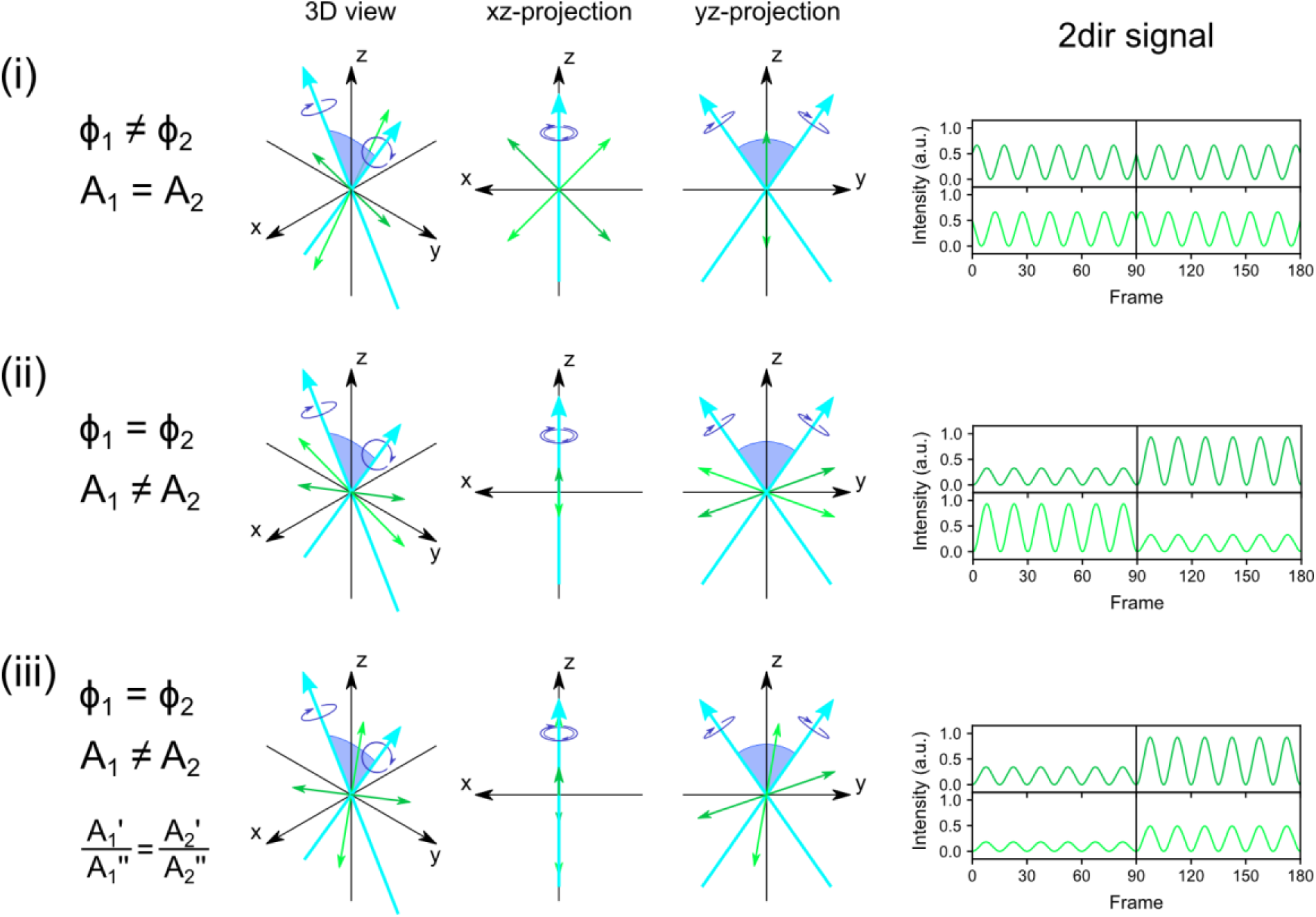
Comparison of the 2dir fluorescence signals produced for two molecules (light and dark green) using the same two illumination directions (light blue). The following special cases are addressed: (i) Both molecules show the same amplitudes due to being symmetrically aligned towards the plane spanned by the illumination directions. They are easily distinguished by their phase information. (ii) Both molecules are coplanar with the illumination directions and therefore have the same phase information for both directions, but they can be distinguished if amplitude information is taken into account. (iii) For any orientation within the plane spanned by the two illumination directions there is another one producing the same signal up to a constant factor. If the absolute brightness of a molecule in this plane is not known, its orientation is therefore ambiguous. The only exception to this is when a molecule is oriented exactly parallel to one illumination direction, since it is the only orientation resulting in zero amplitude for that particular direction.

A third direction linearly independent from the first two will decide any ambiguous case that arises with two projections. Therefore, using three projections (3dir, Fig. 1b) the full three-dimensional orientation of a molecule is uniquely determined. Moreover, without noise the phase information is already enough to reconstruct the underlying orientation, while relative amplitude information can help evaluating noisy data. In any case absolute brightness information and therefore its calibration is not necessary.

On the one hand, three projections uniquely determine an orientation. On the other hand, orientation reconstruction is hampered by data noise. Hence, the question arises if three projections already contain the maximum information on the 3D orientation when taking the limited number of available photons and shot noise in an experiment into account, or if detecting more projections or even continuously changing the direction of projection during the measurement provides more information.

Here, we present a systematic study to explore the differences in the angular accuracy of different wide-field fluorescence microscopy methods that allow measuring three-dimensional molecular orientation using different illumination directions. A method that utilizes a continuous change of the illumination direction is also presented (Fig. 1c) and its performance compared to methods making use of a finite number of projection directions. We used simulated data with exactly known orientations to evaluate angular accuracy of the different methods. We also verified experimentally the applicability of the methods and mathematical models presented in this paper. To ensure comparability of the different methods identical conditions for data acquisition are considered for a given sample regarding illumination intensity, detection efficiency, background and shot noise, and total measurement duration.

## 2. Materials and Methods

### 2.1 Sample preparation

An aqueous solution of Alexa Fluor 488 (10^−8^ mol/L in H_2_O) was diluted 1:100 in a solution of polyvinyl alcohol (PVA, 98-99 % hydrolyzed, low molecular weight, Alfa Aesar; 1.5 % in H_2_O). A droplet (10 µL) of this solution was placed on a microscopy cover slip (borosilicate glass, 0.13-0.16 mm thickness, 22×22 mm, Carl Roth) which was subsequently held sideways and continuously rotated to spread the solution on the surface until all the solvent had evaporated and a thin solid PVA layer containing randomly oriented dye molecules was left.

### 2.2 Experimental setup

The setup used was a self-built wide-field epifluorescence microscope. A 488 nm CW laser beam (Sapphire 488-50, Coherent) was widened by a telescope system (achromatic doublets, f = 30 mm and f = 500 mm, Thorlabs). The peripheral part of the beam was blocked using an iris diaphragm, allowing only the central part (diameter of about 1 cm) to pass through. The beam was reflected by three silver mirrors and then by a polarizing beam splitter cube (PTW 20, 440-650 nm, B. Halle Nachfl. GmbH), before passing a rotating half-wave plate (WPH05M-488, 488 nm, Thorlabs), another lens (achromatic doublet, f = 400 mm, Thorlabs) that was used to focus the light onto the back focal plane of the objective, and two wedge prisms (PS811-A, 4° beam deviation, Thorlabs). The prisms were fixed into a single housing in an antiparallel orientation and with a distance of about 3 cm resulting in a ca. 1.5 mm lateral displacement of the laser beam (cf. Fig. 3). The beam was reflected by a dichroic mirror (XF2045 400-485-580TBDR, Omega Optical) towards the objective (NA = 1.35 oil immersion objective, UPlanSApo, 60x, Olympus) with both mounted into an inverted microscope body (IX71, Olympus). The tilt angle of the beam illuminating the sample was *β* = 35°. The sample’s fluorescence light collected by the same objective passed through the aforementioned dichroic mirror. A mirror redirected it towards the electron-multiplying charge coupled device camera (EMCCD, iXonEM+897 back-illuminated, Andor Technology) which recorded the signal with a frame rate of 30 Hz. Before being detected by the camera the light passed a telescope system (achromatic doublets, f = 40 mm and f = 250 mm, Thorlabs) and two filters (band pass filters, 525/50 and 525/30, AHF) to further enlarge the image and filter out scattered excitation light.

**Fig. 3.**
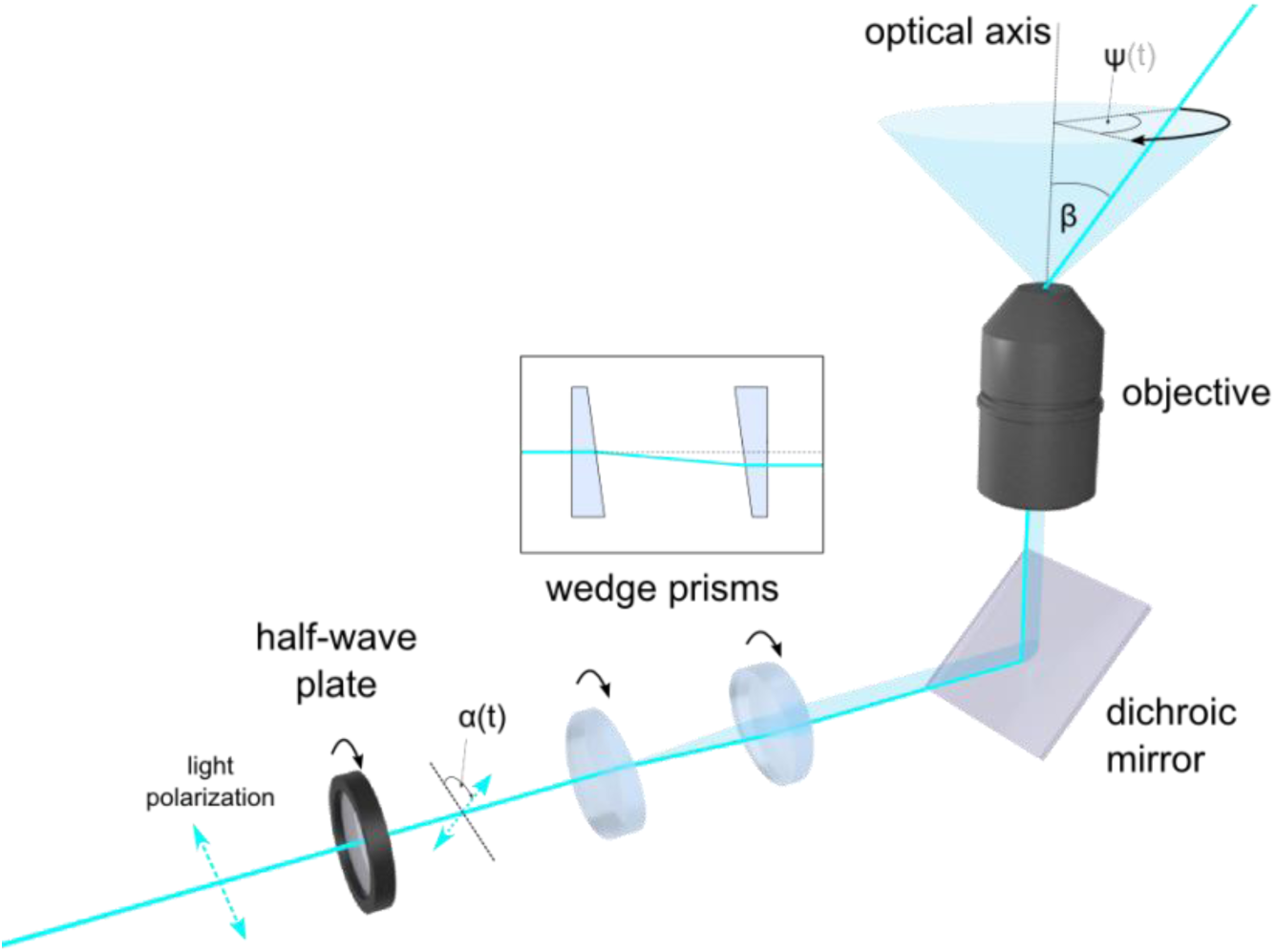
Schematic depiction of experimental setup for the presented multidirectional and double modulation approaches. Linearly polarized light passes a rotating half-wave plate, resulting in a rotation of its polarization plane described by *α*(t). Two wedge prisms in antiparallel orientation laterally displace the light beam. The dichroic mirror (DM) reflects the light onto the objective’s back aperture. Because of the lateral displacement of the beam it exits the objective tilted by a constant angle *β*. Here, for simplicity refraction at the glass-air-interface (after illuminating the sample) is not shown. The tilted beam can be rotated around the optical axis by a rotation of the wedge prisms described by the rotation angle *ψ*. In the multidirectional approach, several measurements with different constant rotational positions are done while in the double modulation approach, the wedge prisms are continuously rotated during a measurement making *ψ* a linear function of time just like *α*.

The rotations of the wedge prisms and the half-wave plate were achieved by connecting the motor of an optical chopper system (OCS, MC2000-FW-SP, Thorlabs) to the respective housing with a rubber belt. The two OCS used the frame-rate of the EMCCD camera as reference for the rotation speed which in turn was controlled using a chopper wheel connected to the housing containing the optical elements.

### 2.3 Measurement procedure

In the double modulation experiments the rotation speed of the two wedge prisms was set to 0.5 Hz, i.e. one full rotation per 60 camera frames. The rotation speed of the half-wave plate was set to 0.5 Hz (*m* = 2), 0.75 Hz (*m* = 3) or 1 Hz (*m* = 4), respectively. For the multidirectional measurements, one OCS was switched off and only the half-wave plate was rotated with a rotation speed of 0.5 Hz. The orientation of the wedge prisms needed for the different illumination directions was set manually before each measurement. The same orientation was used for the first illumination direction in every experiment (*ψ* = 0°). The other orientations were spread evenly over the 360° range, i.e. *ψ* _1_ = 0° and *ψ* _2_ = 180° for two directions and *ψ* _1_ = 0°, *ψ* _2_ = 120° and *ψ* _3_ = 240° for three.

### 2.4 Simulations

The simulations were created using a custom Python script. Per video, a 32 by 32 grid of point emitters with random three-dimensional orientations was generated with their fluorescence signals calculated from Eq. (4) or Eq. (6), depending on the method. A normalized 2D Gaussian distribution with a FWHM of 5 pixels was used as PSF and multiplied by the number of simulated excitation photons (380 per video frame) as well as the orientation dependent detection efficiency (which lies between 18% and 32%). Shot noise was applied to the simulated photons by drawing from a Poisson distribution. We divided the result by 0.041 and then modelled a baseline count and detector noise by adding to each pixel a value drawn from a normal distribution with mean value 100 and standard deviation 30 to translate the photon count into a signal count that we would expect to detect with our camera. All values for brightness and noise levels were chosen to correspond to the values we usually observe in experiments. Four videos using different random orientation sets were created per method. For the multidirectional method, not only simulations for two (2dir) and three (3dir) illumination directions were created, but also for 4dir, 6dir, 8dir, 12dir and 24dir. The angles *ψ* _*i*_ describing the rotational position of the *i*-th illumination direction were always spread evenly over the whole 360° range 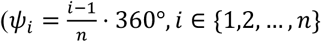 with *n* being the total number of directions. Note that these angle values do not correspond directly to the coordinate system used in Fig. 6, but in comparison are offset by 90°).

**Fig. 4.**
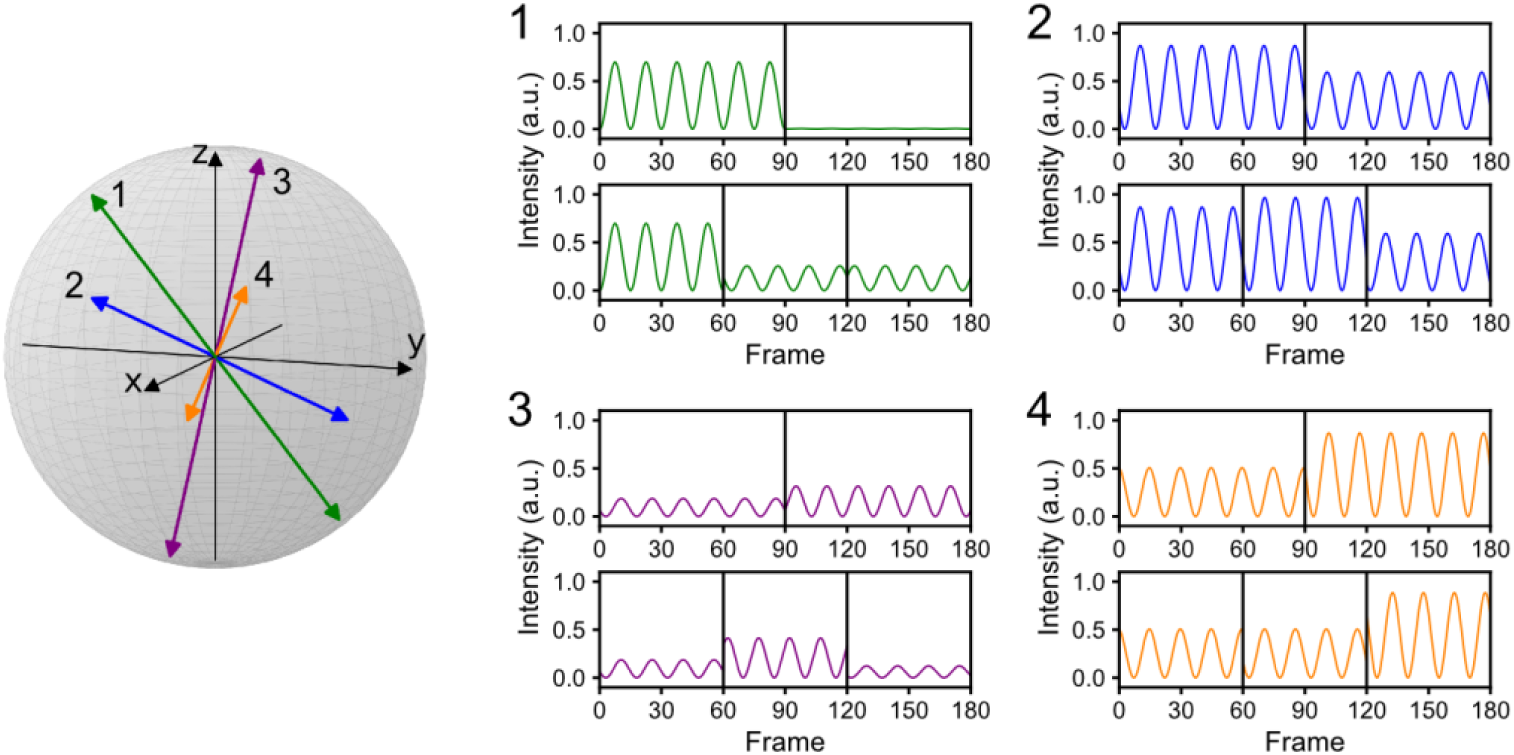
Theoretical example curves of four different orientations observed with the 2dir (1-4, top) and 3dir (1-4, bottom) setup. The orientations are illustrated in a spherical plot and have the following values (*φ*/ *θ* as defined in Eq. (1)): 90°/50° (green), 120°/10° (blue), 200°/70° (violet) and 330°/30° (orange). Orientation dependent detection efficiency is considered in all cases, based on an objective lens with a numerical aperture of 1.35.

**Fig. 5.**
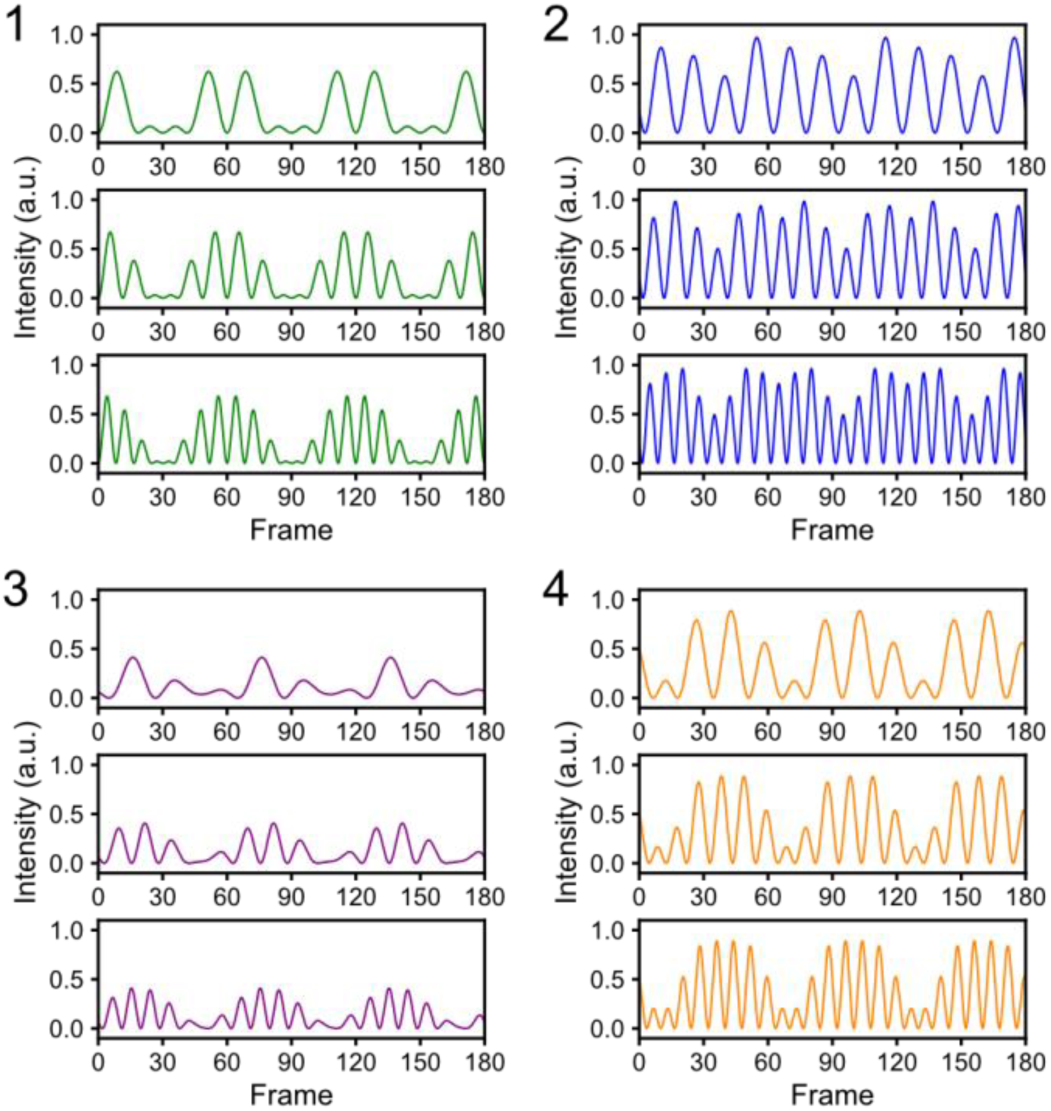
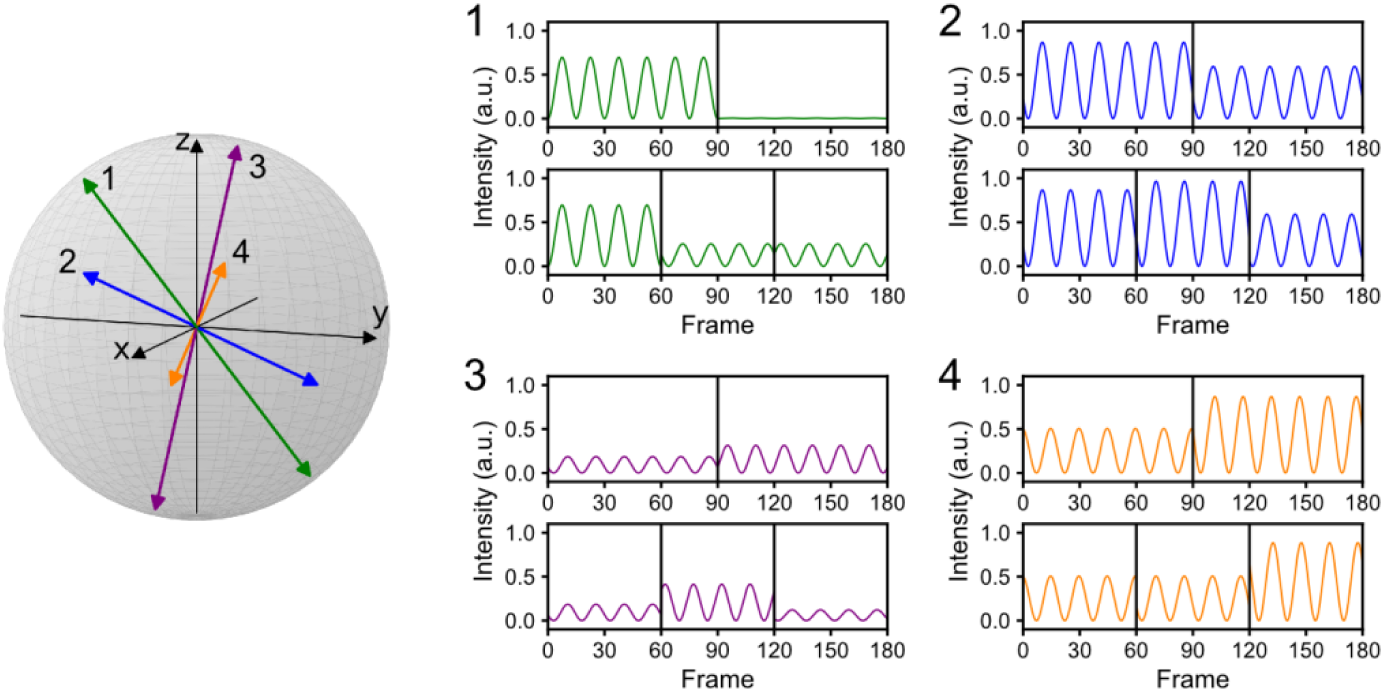
Theoretical example curves of four different orientations observed with the double modulation setup. The same four orientations as in are considered. In each case their fluorescence traces for *m* = 2 (top row), *m* = 3 (middle row) and *m* = 4 (bottom row) are shown, with *m* being the ratio between the rotational velocities of the illumination light’s polarization plane and the illumination direction.

**Fig. 6.**
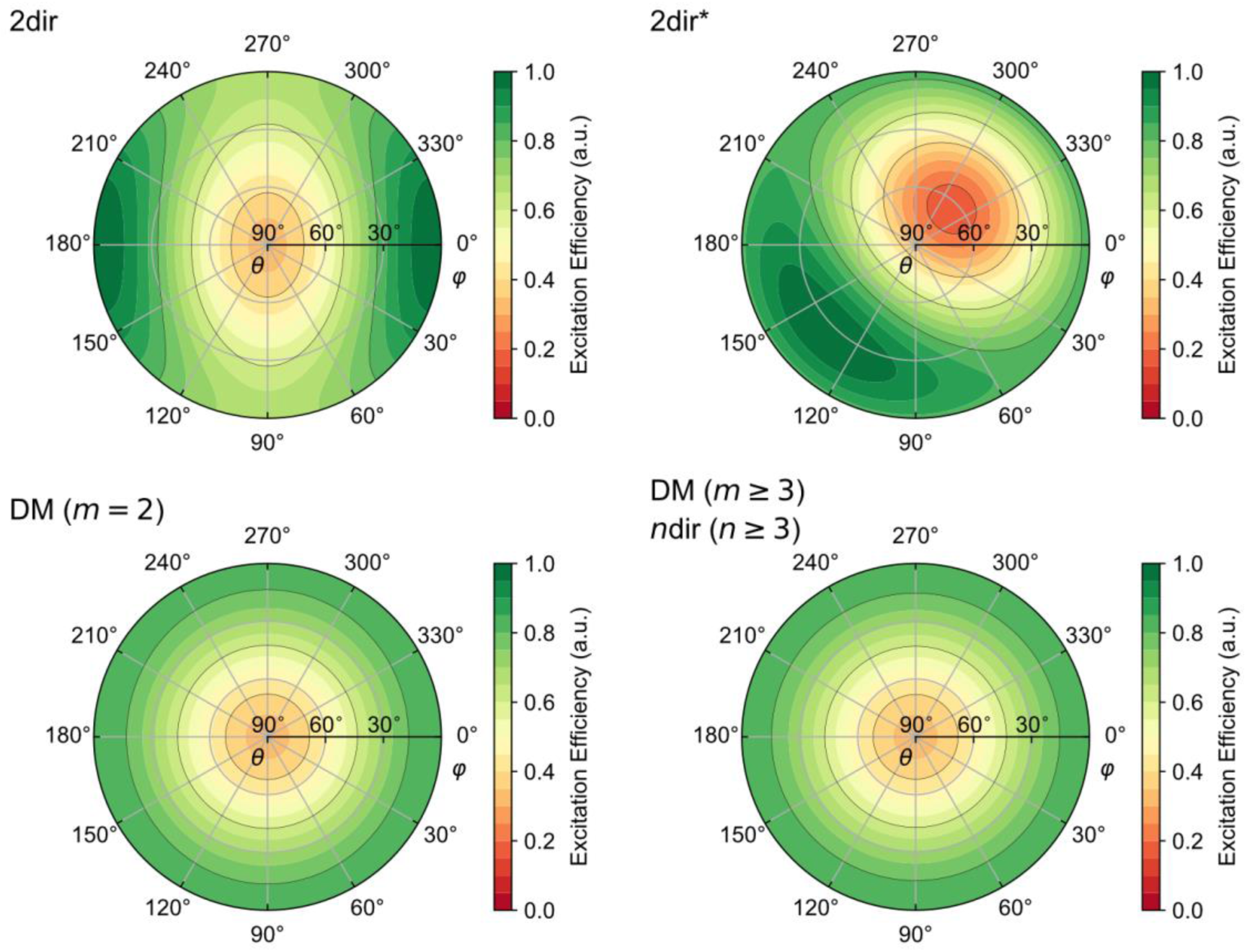
Normalized orientation dependent excitation efficiencies for different polarization modulation methods using *β* = 35°. Maximum efficiency (set to 1) is reached for molecules that are perpendicular to all illumination directions (only possible if two directions are used). In addition to our standard 2dir method (top left, azimuths of illumination directions 90° and 270°, i.e. 180° apart) also shown is the excitation efficiency for two directions with illumination directions 90° apart (top right, 0° and 270°). The comparison shows that further apart directions result in a more homogeneous spatial excitation efficiency. As indicated in the main text, for *β* = 35° almost no difference in excitation efficiency is observed for the double modulation (DM) method using *m* = 2 (bottom left) and all other methods (i.e. double modulation with *m* ≥ 3 and multidirectional methods using three or more directions, bottom right), the excitation efficiencies of which are independent of the molecule’s azimuthal orientation *φ*.

Corresponding simulations were produced with lower excitation rates (190, 85 and 38 excitation photons per frame and molecule) to be able to evaluate the difference in orientation reconstruction accuracy based on photon counts. The detector noise level was not changed.

### 2.5 Theory and Data Analysis

In this manuscript, the molecular orientation is described by spherical coordinate angles. The azimuth angle *φ* is defined here to be 0° when the azimuth is parallel to the x-axis, and growing in a clockwise direction when viewed from above (i.e. from the positive x-towards the negative y-axis). The elevation angle *θ* is used, 0° corresponding to “flat” molecules oriented in the xy-plane and 90° corresponding to molecules oriented parallel to the optical axis (= z-axis). Since a transition dipole moment vector and its negative are not discernible, half of a spherical coordinate system is sufficient to describe all possible orientations. We decided to allow only positive z-components for transition dipole moment vectors. The normalized (unit length) transition dipole moment vector 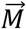 of an emitter is therefore:

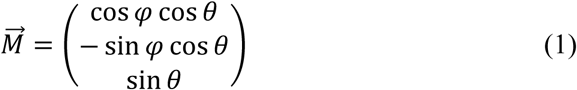

#### Multidirectional method

The behavior of the fluorescence signals observed using multiple distinct directions can be understood quite intuitively as they correspond to “simple 2D-projections” of the transition dipole moment vector. The signals obtained with a certain illumination direction therefore follow a cosine-squared relationship with respect to the orientation of the illumination light’s polarization plane.

Let 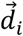 be the unit vector describing the *i*-th direction of illumination and 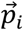 the unit vector in the plane perpendicular to 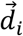 corresponding to phase angle 0°. Furthermore, let 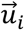 be the projection of an emitter’s normalized transition dipole moment vector 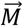 onto said plane, which can be calculated as follows:

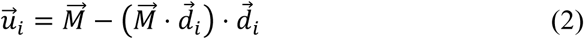

Then, the phase Φ_*i*_ of the cos^2^ fluorescence signal observed with the given illumination direction can be calculated by:

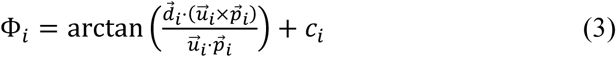

Here, c_*i*_ is a calibration parameter that maps the rotational position of the half-wave plate onto the corresponding video frame. The length of 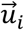 describes the excitation efficiency and is therefore proportional to the fluorescence intensity. Hence, for a given illumination direction *i* the temporal signal *I*_*i*_(t) is described by the following equation (4):

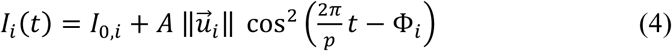

Here, *I*_0,*i*_ and A are offset and amplitude parameters that depend on experimental conditions such as illumination power and camera settings and p is the period length (same unit as time t). In this manuscript we define Φ_*i*_ to be zero when the polarization vector of the illumination light – before being refracted by the objective – is parallel to the sample plane and perpendicular to the first illumination direction (which makes it parallel to the x-axis). In an experiment the corresponding video frame can be found for example by temporarily inserting a polarization filter in the illumination beam path at the beginning of the video.

In Fig. 4, theoretical 2dir and 3dir curves for four exemplary orientations are shown (*φ*/*θ*): 90°/50°, 120°/10°, 200°/70° and 330°/30°. For simplicity, no temporal offset was simulated (i.e. Φ_*i*_ = 0). The smaller the angle between the simulated transition dipole moment and a given illumination direction, the smaller is the overall amplitude of the resulting fluorescence signal. An example for this is molecule 1 which is almost parallel to the second 2dir illumination direction and whose 2dir reconstruction will therefore be inaccurate when noise is present if only phase information is used for evaluation. Orientation dependent detection efficiency is also considered in all the shown graphs, based on an objective lens with a numerical aperture of 1.35. As a consequence, the intensities in example 3 are much lower in comparison to the other examples because of the emitter’s high elevation angle (70°).

#### Double modulation method

We also developed a double modulation scheme where not only the illumination light’s polarization is continuously rotated during a measurement but the direction of incidence is as well. This is achieved by a continuous rotation of the wedge prisms. As a result, the illumination direction rotates around the optical axis while having a constant angle towards it. In our setup the half-wave plate on the one hand and the wedge prisms on the other can be rotated independently from each other. Therefore, different rotational velocity ratios can be realized.

By continuously rotating the illumination direction during a measurement while also modulating the excitation polarization (as illustrated in Fig. 1c), the resulting excitation probability and therefore the fluorescence signal is no longer described by a simple cosine-squared relationship. Rather, the relative excitation probability P can be described the following way, with *β* describing the tilt angle of the illumination direction measured from the objective’s optical axis, and *α* and *ψ* describing the rotation of the light’s polarization and the illumination direction, respectively:

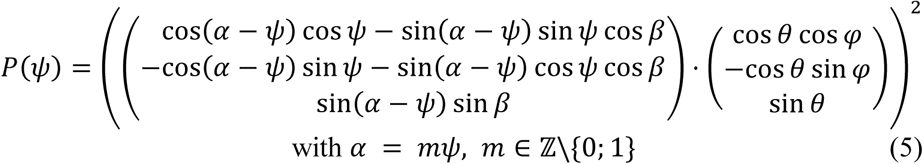

For simplicity, we only looked at integer ratios between the rotational velocities of the two rotations. The ratio is called *m* in this manuscript. For instance, a value of *m* = 2 means the rotation of the light’s polarization plane is twice as fast as the rotation of the illumination direction. An integer ratio keeps the overall period of the signal short, allowing for averaging of several periods. The values *m* = 0 and *m* = 1 are not suitable for orientation measurements as both can lead to ambiguous cases. The values considered in this work are *m* = 2, 3, 4. Theoretically, the signals observed with this setup are described by the following equation:

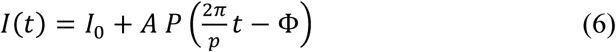

with *P* as defined in Eq. with *α* = *mψ, m* ∈ ℤ\{0; 1} (5) and *I*_0_, *A, p* and Φ having the same meaning as the corresponding parameters in equation (4).

In Fig. 5, theoretical example curves for the double modulation technique with *m* = 2, 3, 4 are shown. The simulated orientations are the same as in Fig. 4. The effects of the two modulations are clearly visible. While polarization modulation causes the expected oscillation between zero intensity and a certain maximum intensity, the latter is limited by the illumination direction modulation. In fact, regardless of the value of *m* the resulting signal is limited by the same envelope curve since the rotation behavior of the wedge prisms is the same. Comparing the different orientations to each other shows that more out-of-plane molecules (see molecule 3) show fewer oscillations per period. Since a parallel orientation of transition dipole moment and illumination direction make excitation impossible, the envelope curve goes down close to zero if the elevation of the molecule is close to the elevation of the illumination direction (see molecule 1).

#### Data Analysis

A simple direct approach to evaluate 2dir, 3dir, etc. data is to use only the phase information that is easily accessible via Fourier Transform of the signals. For each given illumination direction a plane can be determined that contains both the transition dipole moment vector and the illumination direction itself. These planes can be sufficiently described by their normal vectors 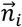 as only orientation and not position is of interest. Necessarily, the transition dipole moment is perpendicular to the normal vectors of all measured directions. In the case of 2dir, the normalized cross product of the two normal vectors 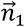 and 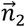 can be used to get a representation for the normalized transition dipole moment vector. The general result of this cross product is shown in the following equation (7), with *α*_1_ and *α*_2_ being the phases obtained from the two directions. This evaluation method was used in a previous manuscript of ours. [17] Note that the expression shown in equation (7) differs from the one shown in that manuscript only by a constant factor; both map to the same vector upon normalization to unity length.

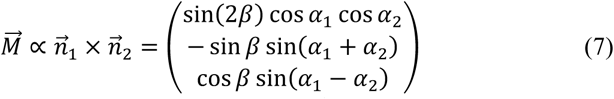

Alternatively, the unknown transition dipole moment vector 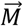 can be found as a non-trivial solution for the homogeneous linear equation system consisting of its dot products with each of the normal vectors set to zero 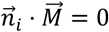 for all directions *i*). If *n* is the total number of illumination directions measured these equations can be summarized into one using the [*n* × 3] matrix **N** that contains all normal vectors as row vectors:

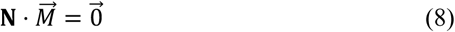

Since real data usually is noisy this equation system generally cannot be expected to be exactly solvable (except for the uninteresting trivial solution 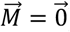. Still, a non-trivial “best” solution can be found numerically via singular value decomposition of N, assuming 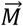 to be the right-singular vector of N that corresponds to the lowest singular value.

Both simulations as well as experimental data were evaluated by first selecting regions of interest (ROIs) containing most of a single emitter’s signal and calculating the average observed intensity for each video frame, resulting in one temporal intensity signal per emitter and video. Depending on the method of measurement, a least-squares fit of Eq. (4) or Eq. (6) to these signals was performed with the orientation parameters *φ* and *θ* obtained as a result, as they are contained in Φ_*i*_ and P(*ψ*), respectively. In the case of simulations the angles between these orientations and the true simulated orientations were used to quantify the reconstruction error. If only phase information was used to evaluate signals produced with the multidirectional method, the signals’ phases were determined via Fast Fourier Transform and translated into normal vectors of planes containing the examined transition dipole moment vector. The molecular orientation was determined via singular value decomposition, see Eq. (8).

#### Excitation Efficiency

In general it can be expected that the more photons are detected from a molecule, the more precise its orientation reconstruction will be. Due to the anisotropic fluorescence intensity profile of fluorophores even with the same excitation rate fewer photons are detected when their transition dipole moment vector has a higher elevation angle. This detection efficiency will be the same for all methods considered in this manuscript as it is only dependent on the detection system and not on the excitation. On the other hand, the absorption behavior of a fluorophore shows the same anisotropy as the fluorescence, so different methods of excitation involving changes in the illumination direction can result in different numbers of detected photons. For a fair comparison, this could in principle be compensated by different excitation intensities, for example. However, the following detailed analysis demonstrates that for a random distribution of molecule orientations the average number of excitations and therefore the number of detected photons is identical in all methods explored here.

The excitation efficiency can be deduced by calculating the integral of the fluorescence signals obtained for a given method and molecular orientation. Fig. 6 shows normalized polar plots corresponding to these integrals using *β* = 35° (calculated using equations (4) and (6) with A = 1, *I*_0_ = 0 and Φ = 0). Here, in addition to the plot for the 2dir method as presented in this manuscript (top left) a plot is shown (top right) for a method using two directions whose azimuths are not opposed to each other but have a 90° angle (*ψ* _2_ ≡ *ψ* _1_ + 90°, as was the case in the method developed by Prummer et al.). This adapted 2dir method is denoted by an asterisk in this manuscript (2dir*). Both of these methods using two directions, the latter even more pronounced, show a rather strong dependence on *φ*, meaning emitters with certain azimuthal orientations are excited more effectively than those with certain other azimuthal orientations. The least effectively excited molecules are those oriented in the middle of the two excitation directions. Therefore, this bias against certain orientations is the stronger the closer to each other the illumination directions are, which is why for *β* < 45° the 2dir method with the directions’ azimuths opposing each other (*ψ* _2_ ≡ *ψ* _1_ + 180°) has the least bias and is therefore the one addressed in the Results section.

The integral for the double modulation technique with *m* = 2 shows only very little dependence on an emitter’s azimuthal orientation *φ* (Fig. 6, bottom left). This is the case in general if *β* is not too large (roughly *β* < 45°). Interestingly, the signals obtained for three or more illumination directions (3dir, 4dir, etc.) result in the same overall integral – taking all directions into account – as the double modulation method for *m* ≥ 3 (Fig. 6, bottom right). This integral is independent of the emitter’s azimuthal orientation *φ*. For example, this concerns the 3dir examples in Fig. 4 and the double modulation examples in Fig. 5 (middle and bottom row) which all have the same area under the curves. This already indicates that at least for noiseless data all these methods should show the same performance in orientation reconstruction as the same number of photons is detected from a molecule with a given orientation. It is important to note that for 3dir, 4dir etc. the same integral is only obtained if the directions are evenly spread over the full 360° range of *ψ* (as previously explained, we used 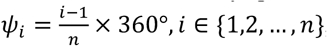, with *n* being the number of directions).

Despite these differences between 2dir, DM (*m* = 2) and the other methods, it is important to note that integrating with respect to *φ* gives the same result for all methods. This means none of the methods is inherently better at exciting molecules, only some methods (especially those using only two directions) can be expected to have preferred orientations that they are better and other orientations that they are worse at reconstructing due to different excitation efficiencies, while the other methods are completely unbiased with respect to azimuthal orientations (circularly symmetric diagram in Fig. 6).

## 3. Results and Discussion

### 3.1 Assessment of the methods’ performance using simulations

To evaluate if the presented methods differ in the accuracy of the determined orientation, we first simulated 4096 randomly oriented molecules and generated the corresponding signals obtained with polarization modulated illumination from 2, 3, 4, 6, 8, 12, and 24 directions that are evenly distributed on a right circular cone around the optical axis (2dir, 3dir, …, 24dir for short). We also generated double modulation signals that are obtained with both the illumination light’s polarization plane and the illumination direction being continuously rotated. Three different rotational velocity ratios were considered, with the polarization plane’s rotation being twice, three times and four times as fast as the illumination direction’s rotation around the optical axis (*m* = 2, *m* = 3 and *m* = 4). To ensure direct comparability we simulated identical numbers of excitation photons (380 photons per frame and molecule) and noise levels (detector noise with a standard deviation equivalent to about 1.2 photons per frame and molecule) as well as the same total number of frames (360). Note that because of both excitation and detection efficiency the number of detected photons per molecule is significantly lower than the number of simulated excitation photons and typically lies between 10 and 40 per frame on average, resulting in an average signal-to-noise-ratio (average signal divided by standard deviation of background noise) between 8 and 33 per frame, or about 150 to 620 for all 360 frames of data, with about 3000 to 15000 detected photons in total.

We then algorithmically determined the orientations of the simulated emitters as explained in section 2.5. The data concerning the multidirectional method was evaluated using only the phase information [Eq. (8)] as well as using all information [fit of Eq. (4)]. As a side note, evaluation of 2dir data using Eq. (7) gives exactly the same results as using Eq. (8) and singular value decomposition. The double modulation data was fit with Eq. (6).

We compared all resulting orientations with the true orientations used for the simulation. Fig. 7a shows a histogram of the obtained orientation reconstruction errors (the absolute acute angle between simulated and reconstructed orientation vector). The corresponding average and median deviations are shown in Fig. 7b. It is immediately obvious that with 2dir the worst reconstruction accuracy is achieved. While the majority of orientations is reconstructed with an accuracy of better than 2°, as is also the case for all other simulations, there is a significant percentage of orientations reconstructed with worse than 10° accuracy. A closer investigation reveals that almost all of these orientations lie within 10° of the plane spanned by the two illumination directions. Surprisingly, fewer of these orientations are reconstructed successfully when considering also the intensity [Eq. (4) fit] than when using only the phase information [Eq. (8)]. Looking at the results for the simulations with more than two directions, using all information always gives better results than using only phase information. For example, about 90 % of orientations being found with 3° accuracy or better using phase information, while this percentage increases to 98 % when using the complete model fit.

**Fig. 7.**
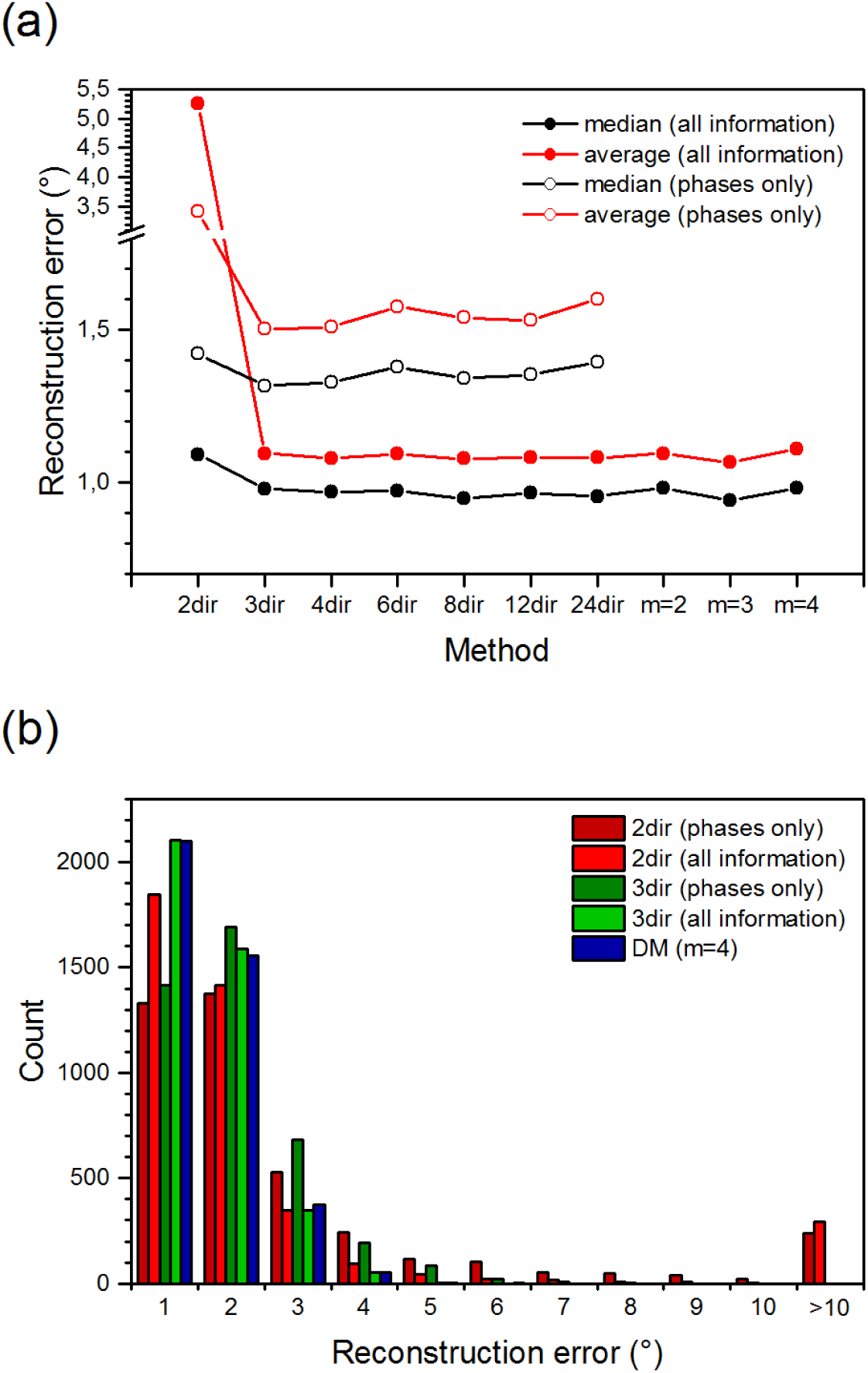
a) Average (red) and median (black) reconstruction errors for all simulations. For the multidirectional simulations the results for both evaluations are shown, i.e. fitting the data with the mathematical model (Eq. (4), full circles) and using only the phases (Eq. (7)–(8), hollow circles). b) Reconstruction errors of different orientation measurement methods applied to simulations of 4096 randomly oriented emitters. 360 frames were evaluated in each case. With all methods the majority of orientations is reconstructed within 2° accuracy. Using only two directions (2dir) a significant percentage of molecules is not reconstructed well at all, with errors larger than 10°. Using three directions (3dir) this is no longer the case. For both 2dir and 3dir using all information gives better results than using only the phase information. The double modulation (DM) method shows similar results to the 3dir evaluation that uses all information.

With an increasing number of directions the phase-only evaluations show a very slight increase in orientation reconstruction error, indicating that the positive effect of knowing more projections of the transition dipole moment is outweighed by the negative effect of a less precise determination of these projections (as fewer frames per direction are evaluated when more directions are used).

When comparing the 3dir fit results to those obtained with more than three illumination directions (4dir etc.) and with the double modulation method, no significant change in accuracy can be observed. It can be concluded, that even with the data distorted by noise three directions already contain the maximum information about the underlying molecular orientation and no information is gained by considering more directions.

Repeating the simulation production and evaluation with lower excitation rates and/or only half of the simulated frames similar results were obtained, with the 2dir results being significantly more inaccurate than the otherwise equally accurate rest of the methods. Table 1 shows the median accuracy obtained for the signals of all 4096 simulated emitters, incorporating the results for all methods except for 2dir. Clearly the accuracy of the results directly corresponds to the signal-to-noise ratio of the evaluated data, as both a higher excitation and thereby fluorescence rate and longer exposure times lead to more accurate orientation reconstructions. This finding can be applied to other mechanisms leading to a different photon count as well, e.g. the use of fluorophores with different fluorescence quantum yields or of objectives with different numerical aperture.

**Table 1.** Median angular accuracies obtained from the simulation evaluations using different excitation intensities and numbers of frames evaluated. All simulations except for 2dir were considered.

| excitation photons per frame and molecule | 180 frames evaluated | 360 frames evaluated |
| --- | --- | --- |
| 38 | 11.3° | 7.9° |
| 85 | 5.2° | 3.7° |
| 190 | 2.5° | 1.8° |
| 380 | 1.4° | 1.0° |

### 3.2 Experimental confirmation of mathematical models

Next, we measured experimental single molecule data with the 2dir, 3dir and double modulation methods (Fig. 8, Fig. 9). Fig. 8 shows the average wide-field microscopy image of the evaluated videos. Additionally, enlarged crops of three molecules are shown in close-up with enhanced contrast. Fig. 9 shows the measured data for these three selected molecules together with their fit evaluations. Interestingly, the steep out-of-plane orientation of molecule III (*θ* ≈ 70°) that is obtained from the fits is also visible in the raw image, as its fluorescence signal shows a ring pattern, which is typical for steep molecules when they are slightly out of focus. [3] The orientations obtained from the evaluations are shown in Table 2. Since the angular difference between three-dimensional orientations is hard to deduce from the spherical coordinates alone, as a measurement of similarity between the results of the different methods we use the maximum angular difference that is obtained when comparing any two of the obtained orientations. All five applied methods are in good agreement with each other, with all results lying within a few degrees of each other (maximum angular differences of 13.6°, 11.3° and 12.0° in the shown examples). The maximum angular differences being lower when the 2dir result is excluded (3.9°, 10.3°, 9.6°) shows that it is generally the least precise method. Two reasons can be given for the particularly strong deviation of the 2dir result of molecule I from the results obtained with other methods. First, the molecule is oriented closely to the second illumination direction, resulting in a very low signal for which phase determination is imprecise. Second, as previously explained, orientations close to the plane spanned by the two illumination directions are not uniquely determinable with 2dir even with correct phase information (e.g. with *β* = 35° the orientations 90°/40° and 90°/68° will result in the same phase and relative amplitude information). In the case of exemplary molecule I the inaccuracy is amplified by the fact that the molecule is apparently oriented along the second illumination direction resulting in a very low fluorescence signal amplitude (see corresponding graph in Fig. 9). Here, data noise leads to a wrong reconstruction of the amplitude which leads to a false reconstruction of the molecule’s elevation angle.

**Table 2.** Orientations obtained from the data shown in Fehler! Verweisquelle konnte nicht gefunden werden. (φ/θ). In the sixth row the maximum angular difference that is obtained when comparing the orientations pairwise. In the last row, the same quantity is shown with the 2dir result excluded.

| Method | Molecule I | Molecule II | Molecule III |
| --- | --- | --- | --- |
| 2dir | $88.9^\circ / 39.6^\circ$ | $349.3^\circ / 6.9^\circ$ | $186.5^\circ / 65.4^\circ$ |
| 3dir | $90.5^\circ / 53.2^\circ$ | $355.3^\circ / 7.5^\circ$ | $209.7^\circ / 68.2^\circ$ |
| DM (m=2) | 87.8° / 49.7° | 349.8° / 16.4° | 200.0° / 63.9° |
| DM (m=3) | 91.3° / 49.8° | 357.2° / 9.8° | 213.6° / 71.7° |
| DM (m=4) | 86.6° / 52.2° | 357.1° / 15.3° | 214.6° / 71.9° |
| Maximum angular difference | 13.6° | 11.3° | 12.0° |
| Maximum angular difference<br>(without 2dir) | 3.9° | 10.3° | 9.6° |

**Fig. 8.**
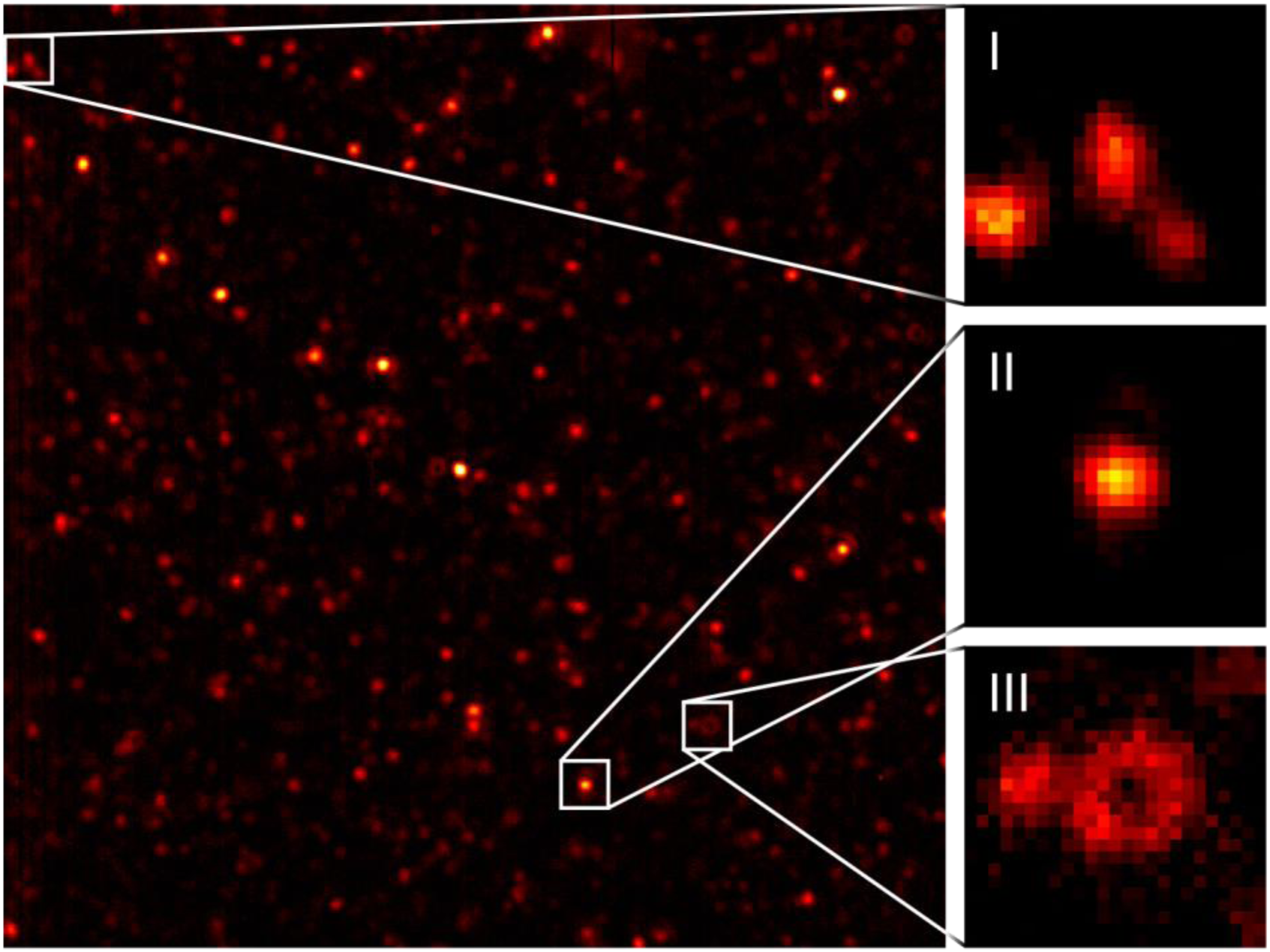
Wide-field microscopy image of single molecules. The insets show enlarged crops with improved contrast containing the single molecules whose data is shown in Fig. 9.

**Fig. 9.**
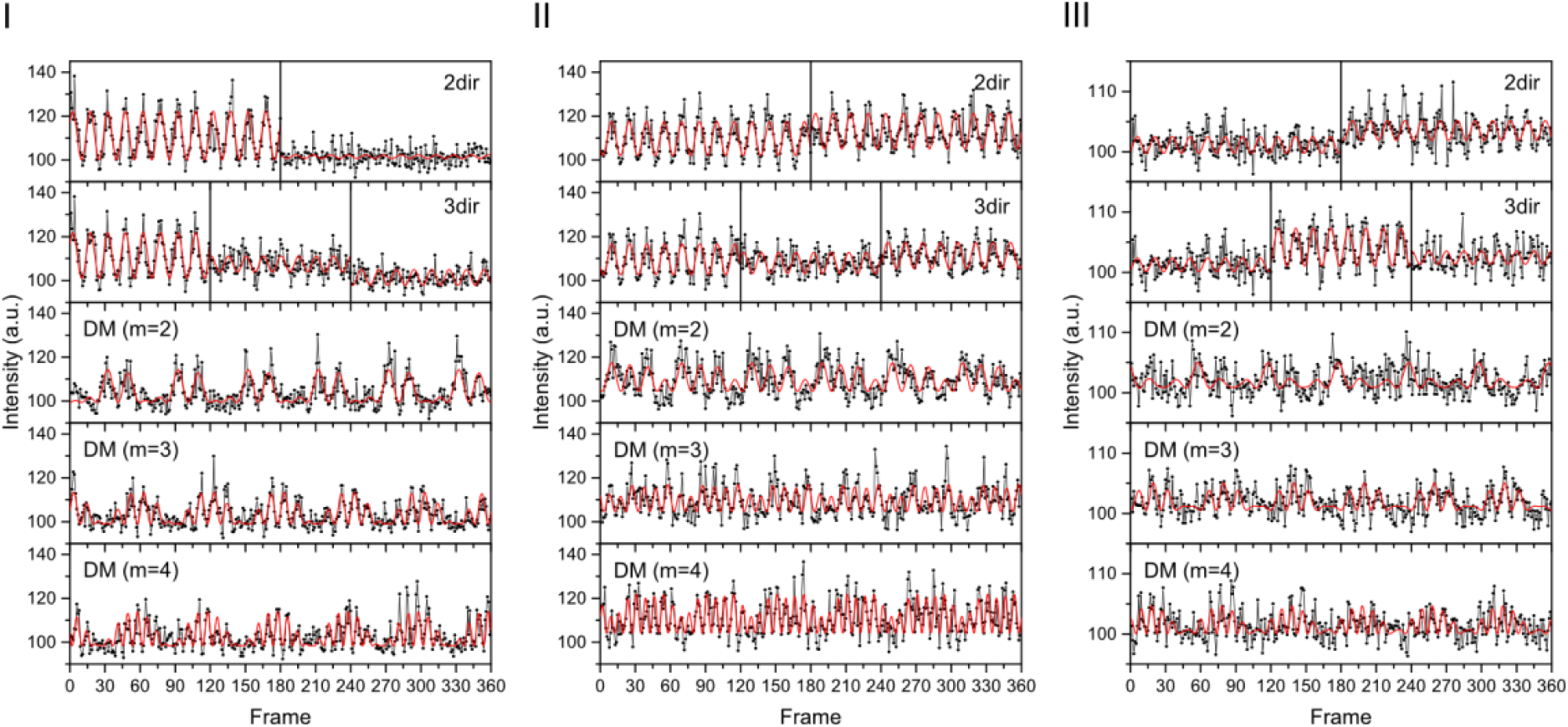
Exemplary 2dir, 3dir and double modulation signals observed in experiments (cf. Fig. 8) together with their fits (see Eq. (4) and (6)). Taking time calibration offsets into account, the average orientations recovered from the different methods are 89°/49°, 354°/11°, 204°/69° (φ/θ). The exact results of all five methods are shown in Table 2.

Compared to the experimental data, the simulated data show better accuracy of as well as agreement between the different methods. This is partly due to the fact that for the simulations blinking of dyes was not considered. Moreover, as multiple measurements with the same sample molecules needed to be conducted the illumination power in the experiment was set lower than usual to reduce the probability of photobleaching, thereby deteriorating the signal-to-noise ratio. Also, slight rotational mobility of the fluorophores might be present causing different results, while the simulations used spatially fixed emitters for all methods. Still, the experimental results clearly corroborate the applicability of the mathematical models presented in this manuscript as a means to molecular orientation reconstruction.

## 4 Conclusion

In summary, we examined different methods to measure the three-dimensional orientation of fluorescence dyes using fluorescence microscopy based on illumination with modulated linearly polarized light from different directions. We investigated the use of different numbers of illumination directions (from 2 up to 24) as well as a double modulation technique that uses a continuously changing illumination direction and polarization modulation with identical or different frequencies. Our results indicate that even with data noise three distinct illumination directions suffice to obtain the maximum information about the orientation of a dye molecule. Using more directions or continuously rotating the direction both do not result in a significant change in orientation reconstruction accuracy. Still, the accuracy can be improved by increasing the signal-to-noise ratio, e.g. longer exposure times. The practical applicability of the different methods (2 directions, 3 directions and three double modulation methods) was experimentally verified.

## Acknowledgements

This work was supported by generous grants from the German science foundation (Deutsche Forschungsgemeinschaft (DFG) (INST 188/334-1 FUGG, GRK2223))

